# Intact mast cell content during mild head injury is required for the development of latent pain sensitization - implications for mechanisms underlying post-traumatic headache

**DOI:** 10.1101/392928

**Authors:** Dara Bree, Dan Levy

## Abstract

Posttraumatic headache (PTH) is one of the most common, debilitating and difficult symptoms to manage after a traumatic head injury. While the mechanisms underlying PTH remain elusive, recent studies in rodent models suggest the potential involvement of calcitonin gene-related peptide (CGRP), a mediator of neurogenic inflammation, and the ensuing activation of meningeal mast cells (MCs), pro-algesic resident immune cells that can lead to the activation of the headache pain pathway. Here, we investigated the relative contribution of MCs to the development of PTH-like pain behaviors in a model of mild closed head injury (mCHI) in male rats. We initially tested the relative contribution of peripheral CGRP signaling to the activation of meningeal MCs following mCHI using an anti-CGRP monoclonal antibody. We then employed a prophylactic MC granule depletion approach to address the hypotheses that intact meningeal MC granule content is necessary for the development of PTH-related pain-like behaviors. The data suggest that following mCHI, ongoing activation of meningeal MCs is not mediated by peripheral CGRP signaling, and does not contribute to development of the mCHI-evoked cephalic mechanical pain hypersensitivity. Our data, however, also reveals that the development of latent sensitization, manifested as persistent hypersensitivity upon the recovery from mCHI-evoked acute cranial hyperalgesia to the headache trigger glyceryl trinitrate requires intact MC content during and immediately after mCHI. Collectively, our data implicate the acute activation of meningeal MCs as mediator of chronic pain hypersensitivity following a concussion or mCHI. Targeting MCs may be explored for early prophylactic treatment of PTH.

## 1. Introduction

One of the most common and disabling symptoms of traumatic head injury is post-traumatic headache (PTH). The estimated prevalence of PTH ranges between 30% to 90% [25,45]. Chronic PTH (lasting more than 3 months) has been reported in about 40% of individuals diagnosed with brain injury attributed to head trauma [9]. While the symptom profiles of individuals suffering from PTH vary, it most often resembles the most common types of primary headaches, namely migraine or tension-type headache [21,24]. The resemblances of PTH to primary headaches points to the possibility that the two conditions are mediated by similar, or shared mechanisms.

A large body of preclinical evidence now supports the view that primary headaches, and particularly migraine, involve the activation and sensitization of trigeminal primary afferent neurons that innervate the intracranial meninges [16,27,32]. Persistent activation of meningeal afferents leading to the sensitization of second-order trigeminal dorsal horn neurons, which receive convergent sensory input from the meninges and cephalic skin has been reported in preclinical models and is thought to underlie the cephalic pain hypersensitivity in primary headaches [31]. The finding of cephalic mechanical hypersensitivity also in PTH [3,4] as well as in preclinical models of mild traumatic brain injury (mTBI) involving a direct cortical impact [26] or a closed-head injury [2,30,44] further supports the notion that PTH and primary headache involve a shared mechanism.

The endogenous processes responsible for driving meningeal primary afferent neurons in primary headache and potentially also in PTH remain poorly understood. One key hypothesis implicates sterile meningeal inflammation, involving the local activation of proinflammatory resident immune cells such as mast cells (MCs) by local release of sensory neuropeptides from meningeal afferent nerve endings via an axon-reflex [15,20,40]. Acute activation of meningeal MCs leading to their degranulation (the release of preformed granule-associated mediators) and local action of their numerous pro-inflammatory mediators have been shown to promote the activation and sensitization of meningeal afferents and the headache pain pathway [17,50]. Degranulation of meningeal MCs can also lead to the development of cephalic mechanical pain hypersensitivity [19]. Our previous finding of persistent degranulation of meningeal MCs in a mouse model of mild closed head injury (mCHI) [18] has led us to hypothesize that such head trauma-related meningeal inflammation is also involved in mediating PTH. The earlier reports that CGRP can promote meningeal MC degranulation [34,39] and that peripheral CGRP signaling mediates PTH-like pain behaviors [2] further points to a potential link between CGRP, meningeal MCs, and the development of PTH.

The aim of the following study thus was to investigate the relative contribution of activated MCs to the development PTH-like pain behaviors using our recently developed rat model of mCHI [2]. We initially investigated the relative contribution of peripheral CGRP signaling to the persistent activation of meningeal MCs following mCHI, using a blocking antibody that selectively targets peripheral CGRP signaling. We then tested the hypothesis that intact meningeal MC granule content during the induction of mCHI is necessary for the development of PTH-related acute and persistent pain-like behaviors following mCHI.

## 2. Material and Methods

### 2.1 Animals

All experiments were approved by the institutional Animal Care and Use Committee of the Beth Israel Deaconess Medical Centre, and were in compliance with the ARRIVE (Animal Research: Reporting of *in vivo* Experiments) guidelines [13]. While females do not appear to be at increased risk of PTH over males [23], given the higher incidence of head injuries in males [41] and that the bulk of clinical data related to PTH primarily reflect finding in males (particularly in military population), we focused on studying male rats. Animals (Sprague-Dawley rats, Taconic, USA, weighing 220–250 g at time of arrival) were housed in pairs with food and water available *ad libitum* under a constant 12 hour light/dark cycle (lights on at 7:00 am) at room temperature. All procedures and testing were conducted during the light phase of the cycle (9:00 am to 4:00 pm). Experimental animals (n = 8-12 per group) were randomly assigned to either sham or mCHI as well as to the different treatment groups.

### 2.2 Experimental mild closed-head impact injury (mCHI)

mCHI was induced using a weight-drop device as described previously [2]. Briefly, rats were anesthetized with 3% isoflurane and placed chest down directly under a weight-drop concussive head trauma device. The device consisted of a hollow cylindrical tube (inner diameter 2.54 cm) placed vertically over the rat’s head. To induce a head trauma, a 250 g projectile was dropped through the tube from a height of 80 cm, striking the center of the head. To ensure consistency of the hit location, animals were placed under the weight drop apparatus so that the weight struck the scalp slightly anterior to the center point between the ears. A foam sponge (thickness 3.81 cm, density 1.1 g/cm^3^) was placed under the animals to support the head while allowing some linear anterior-posterior motion without any angular rotational movement at the moment of impact. A repeated strike was prevented by capturing the weight after the first strike. All animals regained their righting reflex within 2 min (which likely reflect the recovery from anesthesia). Immediately after the impact, animals were returned to their home cages for recovery and were neurologically assessed in the early hours and days post-injury for any behavioral abnormalities suggestive of a major neurological (i.e. motor) impairment. Sham animals were anesthetized for the same period of time as mCHI animals, but not subjected to the weight drop There was 0% mortality after the mCHI procedure with no evidence of skull fractures or cortical bleeding. In this rat mCHI model, animals do no show changes in total distance moved in open field [2] suggesting lack of motor impairment. However, mCHI animals display reduced vertical rearing behavior [2], as in other head injury models, indicating deficits in exploratory behavior likely due to mild traumatic brain injury [8,12].

### 2.3 Prophylactic MC granule depletion protocol

Repeated administration of escalating doses of the MC secretagogue compound 48/80 leads to depletion of connective tissue MC granules with ensuing reduction of MC mediators content that undergoes a slow recovery over several weeks [7,10]. A modified 48/80 injection protocol, used in our previous work [49], was employed to deplete meningeal MC granules prior to mCHI. Briefly, rats were pretreated i.p. with compound 48/80 (0.1% w/v in sterile saline; Sigma-Aldrich) twice a day (morning and afternoon) for a total of eight doses, starting with 0.5 mg/kg injections for the first day, 1 mg/kg for the second, and 2, and 4 mg/kg for the third, and fourth days respectively. To minimize the inflammatory and potentially afferent sensitizing effects elicited by the 48/80-induced MC degranulation, mCHI was induced 4 days following the last 48/80 dosing. In control experiments, animals were subject to a similar injection protocol using the 48/80 vehicle (saline).

### 2.4 Anti-CGRP treatment

Blockade of peripheral CGRP signaling was achieved by using a murine anti-CGRP mAb. Control treatment was a corresponding isotype IgG. Both agents were provided by TEVA Pharmaceuticals, and formulated in phosphate buffered saline (PBS). Anti-CGRP mAb and IgG were injected i.p at a dose of 30 mg/kg, immediately after the head injury and then again 6 days later. This dose has been shown previously to alleviate PTH-like pain behavior (i.e. cephalic allodynia) following mCHI in rats [2] and pain behaviors in other chronic migraine models [14].

### 2.5 Tissue preparation, immunohistochemistry and quantitative assessment of meningeal MC density and degranulation

Animals were deeply anaesthetized with urethane (1.5 g/kg, i.p.) and perfused transcardially with 200 ml of heparinized PBS followed by 150 ml of 4% paraformaldehyde in PBS. The heads were post-fixed overnight in the same fixative solution and then transferred to PBS. Following removal of the calvaria, the intracranial dura was removed bilaterally and mounted on a glass slide. For histological assessment of meningeal MCs, fixed dural whole-mount tissues were stained with toluidine blue (TB, 0.1% in 2.5 pH sodium chloride, Sigma-Aldrich), which binds to glycosaminoglycans in connective tissue MC granules. TB-stained MCs were observed using bright-field illumination, under a 400X magnification (Eclipse Ci, Nikon, Tokyo, Japan). Because MC degranulation levels were uniform across the dura on each side, MC counts and degranulation levels on each side were averaged based on 20 different randomly chosen visual fields. MCs were considered acutely degranulated if there was an extensive dispersion of more than 15 extruded granules localized near the cell, or when there was an extensive loss of granule staining, giving the cell a ‘ghostly’ appearance [18]. Following 48/80 treatment, we also designated irregular-shaped MCs as cells at various stages of recovery, such as cells with abnormal shape (mostly smaller), or with reduced number of TB-stained granules. It is important to note that MCs with fully depleted granules (i.e. fully degranulated) MCs, as expected following repeated 48/80 treatments, cannot be visualized using TB staining, or any other conventional histological approach that is employed to detect MCs and thus were not included in histological analysis. Histological examination of MC staining was conducted by two investigators, and the level of degranulation and presence of abnormal cells were assessed in a blinded fashion.

### 2.6 Assessment of cephalic tactile pain hypersensitivity following mCHI

Behavioral testing were performed during the light phase (09:00-15:00) using a method previously used by us and others to study PTH- and migraine-related pain behaviors [2,6,33,47]. Briefly, animals were placed in a transparent flat-bottomed acrylic holding apparatus (20.4 cm x 8.5 cm). The apparatus was large enough to enable the animals to escape the stimulus. Animals were habituated to the arena for 15 minutes prior to the initial testing. To determine if animals developed pericranial tactile hypersensitivity (i.e. mechanical allodynia) following mCHI, the skin region, including the midline area above the eyes and 2 cm posterior, was stimulated with different von Frey (VF) filaments (0.6 g-10 g) (18011 Semmes-Weinstein Anesthesiometer Kit). Although skin swelling or hematoma were never observed following the trauma, we tested for pain hypersensitivity rostral to the weight drop site to limit the confounding effect of possible local changes related to skin trauma. During the acute phase (3-14 days post-mCHI), we evaluated changes in withdrawal thresholds, as well as non-reflexive pain responses to stimulation using a method previously described [19,51] by recording 4 behavioral responses adapted from Vos et al. [46]. These behavioral responses included: 0) *No response*: rat did not display any response to stimulation 1) *Detection*: rat turned its head towards the stimulating object and explored it, usually by sniffing; 2) *Withdrawal*: rat turned its head away or pulled it briskly away from the stimulating object (which usually followed by scratching or grooming of stimulated region); 3) *Escape/Attack*: rat turned its body briskly in the holding apparatus in order to escape the stimulation or attacked (biting and grabbing movements) the stimulating object. Starting with the lowest force, each filament was applied 3 times with an intra-application interval of 5 seconds and the behavior that was observed at least twice was recorded. For statistical analysis, the score recorded was based on the most aversive behavior noted. The force that elicited three consecutive withdrawal responses was considered as threshold. To evaluate pain behavior in addition to changes in threshold, for each rat, at each time point, a cumulative response score was determined by combining the individual scores (0–3) for each one of the VF filaments tested. Animals were assigned to the different groups in random and all tests were conducted and evaluated in a blinded manner. Responses to VF test stimuli were tested at baseline prior to mCHI, and then at 3, 7, and 14 days post-mCHI.

### 2.7 Latent mechanical sensitization following mCHI

The development of cephalic and hind paw latent mechanical hyperalgesia in response to systemic administration of a previously subthreshold dose of the headache trigger glyceryl trinitrate (GTN; 100μg/kg i.p., American Reagents, USA) was assessed as described [2]. Briefly, rats depleted of their MC granules pre-mCHI, using the 48/80 protocol, and control rats receiving only vehicle were subjected to testing of cephalic mechanical pain thresholds at day 29 post mCHI (pre-GTN baseline) as described above. Baseline hind paw mechanical sensitivity was also tested by assessing withdrawal responses following mechanical VF stimulation of the mid-dorsal part of the hind paw. On the next day, the animals received GTN and were assessed for changes in cephalic and hind paw mechanical pain thresholds 4 hours later.

### 2.8 Data analyses

Data are presented as mean + standard error of the mean. Statistical analyses were conducted using GraphPad Prism (version 7.0). Mechanical threshold (von Frey) data was log-transformed, which is more consistent with a normal distribution [29]. All data passed the normality and homogeneity of variance tests (Shapiro-Wilk and Levene tests, respectively) and therefore was subject to parametric statistic tests. To determine the relative contribution of CGRP to the mCHI-evoked MC responses we employed one-way ANOVA followed by Fisher’s LSD post hoc tests. Correction for multiple comparisons (Type I error) was conducted using the Benjamini and Hochberg false discovery rate (FDR) controlling procedures [1], with Q value set as 0.1. The effects of 48/80 and GTN on MC responses was assessed using unpaired student’s t-test. Differences in pain behaviors between control and 48/80-treated animals at the various time points following mCHI were analyzed using a mixed-design ANOVA to determine the effects of time and treatment. To correct for violation of sphericity, we applied the Greenhouse-Geisser procedure. Post-hoc tests and FDR corrections were as above. The development of latent sensitization to GTN was analyzed using paired, two-tailed student’s t-test. P values, and FDR-corrected p values (q-values) < 0.05 were considered statistically significant.

## 3. Results

### 3.1 mCHI-induced meningeal MC response does not involve peripheral CGRP signaling

We first asked whether blocking peripheral CGRP signaling, using a regimen of treatment with an anti-CGRP mAb that ameliorates mCHI-evoked PTH-like pain hypersensitivity [2] might inhibit the persistent degranulation of meningeal MCs in response to mCHI as reported previously [2]. To test the relative contribution of CGRP signaling in mediating this mCHI-evoked meningeal MC response, and its potential contribution to PTH, we focused on day 7 post-mCHI because it was the earliest time point that post-mCHI anti-CGRP mAb treatment had an anti-hyperalgesic effect in this model [2]. One-way ANOVA revealed a significant difference between the sham and the different mCHI treatment groups on the level of meningeal MC degranulation (F(3,38) = 13.3; p < 0.0001; **Figure 1**). Post-hoc analyses indicated that administration of the anti-CGRP mAb, immediately post mCHI, and again 6 days later, did not decrease the level of meningeal MC degranulation at 7 days post mCHI, which was significantly higher than that observed in untreated animals following a sham mCHI (q < 0.0001), but not different than that observed at the same time point in untreated mCHI animals, or IgG-treated mCHI animals (q = 0.54 for both comparisons). This data suggests therefore that peripheral CGRP signaling does not mediate the ongoing meningeal MC degranulation in this mCHI model.

**Figure 1:**
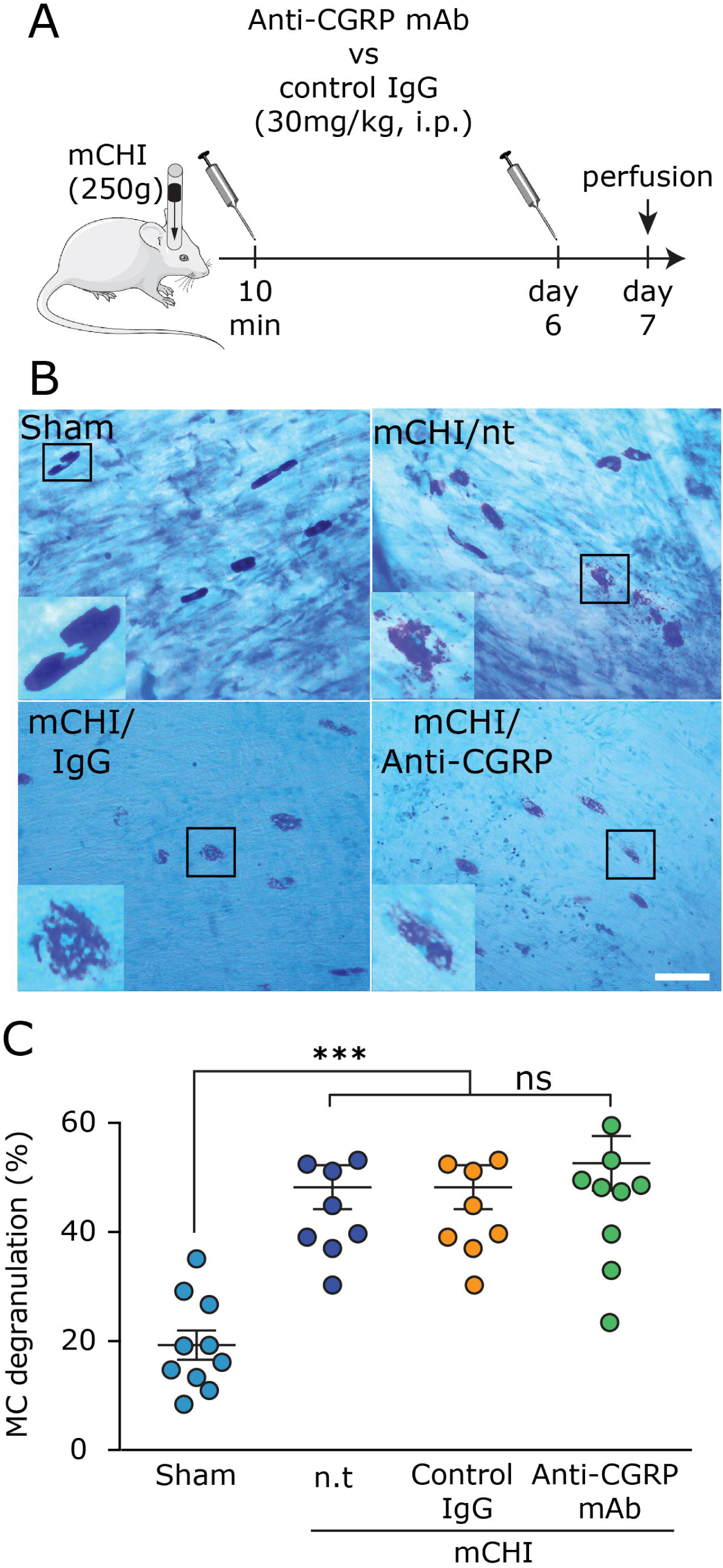
Blockade of peripheral CGRP signaling using systemic administration of an anti-CGRP mAb does not inhibit mCHI-evoked persistent meningeal MC degranulation. (**A)** Scheme of the experimental design. Rats were injected i.p. with anti-CGRP mAb, or control IgG, immediately following mCHI and then 6 days later followed by perfusion a day later (day 7 post mCHI). (**B)** Representative images of TB-stained meningeal whole-mounts showing non-degranulated MCs in animals 7 days following a sham procedure (control) and degranulated MCs in treated, and non-treated (n.t) mCHI animals. Scale bar = 50 μm. (**C)** Comparison of meningeal MC degranulation showing increased levels in 7 days mCHI animals that received no treatment, were treated with the anti-CGRP mAb, or its control IgG when compared to untreated animals undergoing a sham procedure 7 days earlier. *** q < 0.0001 vs sham (FDR corrected values following Fisher’s LSD post-hoc test).

### 3.2 Functional meningeal MCs are not required for development of acute cephalic mechanical hypersensitivity following mCHI

While our data suggests that CGRP contributes to pain hypersensitivity following mCHI via a mechanism unrelated to the ongoing degranulation of meningeal MCs, it does not rule out the possibility that CGRP acts downstream, or in parallel to a MC related process: that the acute MC response post-mCHI provides sufficient meningeal nociceptive stimulus that contributes to the development of PTH-like cephalic pain hypersensitivity, but independent of CGRP signaling. We therefore asked whether limiting the acute mCHI-evoked meningeal MC response, using a prophylactic depletion of their content, might affect the development of the mCHI-evoked cephalic pain hypersensitivity. Meningeal MC granule content was depleted using a 4 day treatment protocol with ascending doses of compound 48/80 as described in the Methods section. Histological examination revealed that the 48/80 treatment protocol was effective in depleting meningeal MC granules at the time of mCHI induction (day 0 in **Figures 2B & C**). At this time point, the total number of non-degranulated meningeal MCs observed in 48/80-treated rats was reduced by 84.6% when compared to that observed in saline-treated animals (p < 0.0001; **Figure 2F**). Very few of the remaining TB-stained MCs were fully granulated and most were comprised of irregular-shaped cells (see example in Figure 2F), likely representing cells at various stages of recovery. At 3 days post mCHI (**Figures 2D, E & G**), the MC population in the control group, was primarily comprised of fully granulated and degranulated cells, while in the 48/80-treated group we observed primarily MCs with irregular shape and almost no fully granulated cells (p < 0.001 vs. the control group, for both comparisons). In this group, we also observed reduced number of cells showing signs of ongoing degranulation (p < 0.001 vs. the control group).

**Figure 2:**
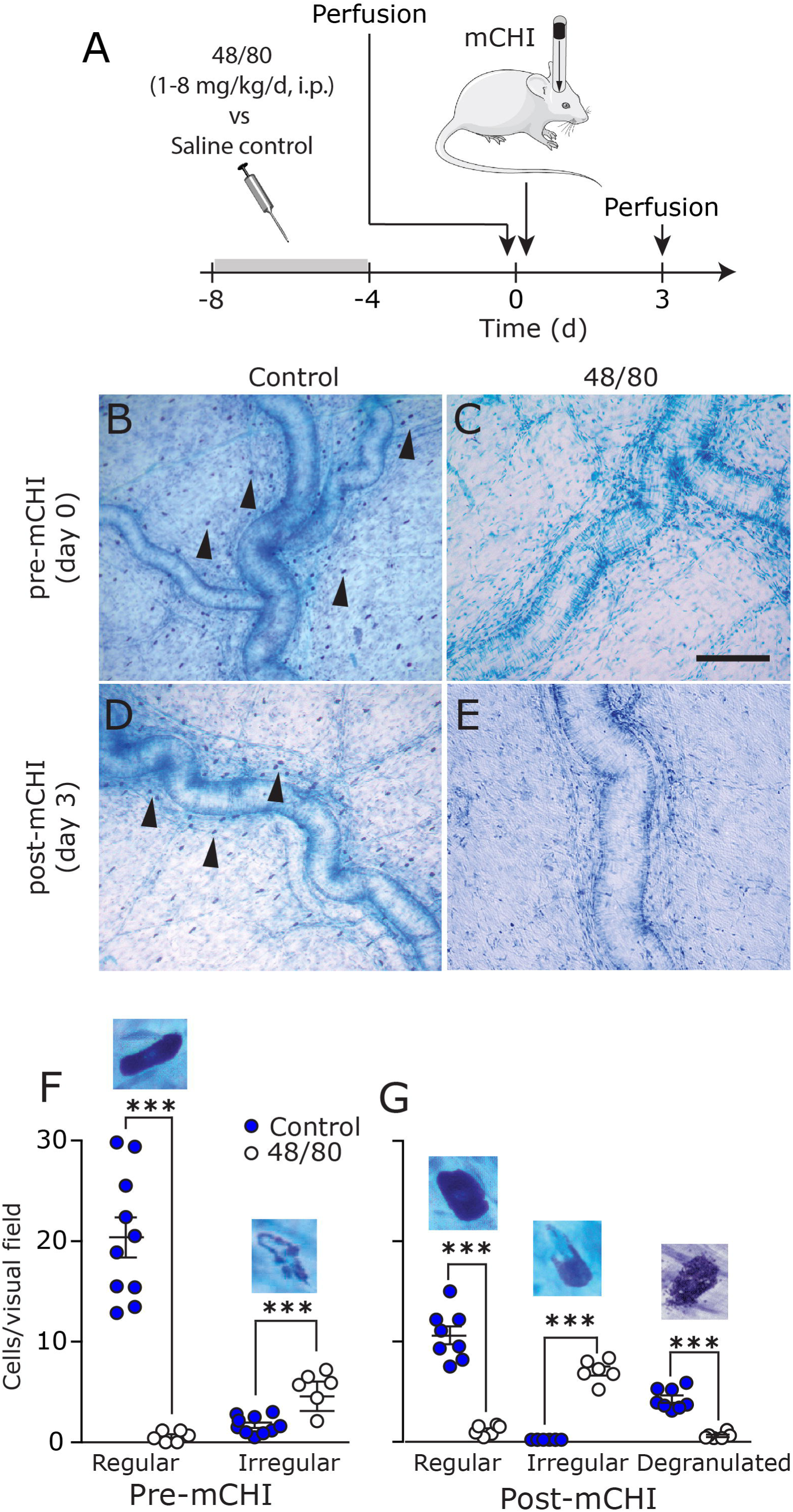
Prophylactic depletion of meningeal MCs using 48/80 treatment. (**A)** Scheme of the experimental design. After 4 days of 48/80 injections, rats were perfused prior to mCHI (day 0), or 3 days post-mCHI. (**B)** Representative examples of TB-stained meningeal whole-mounts depicting numerous MCs (arrow heads) near the middle meningeal artery in animals subjected to control treatment. (**C)** Lack of MC staining in 48/80 pre-treated animals prior to mCHI. Scale bar 100 μm. At 3 days post mCHI, there are numerous MCs in saline pretreated animals showing signs of degranulation (D), but very few MCs in 48/80-treated animals (**E)**. (**F-G)** Quantification of meningeal MCs density in control and 48/80-treated animals. Also shown are high magnification examples of non-degranulated MC in saline treated animals, irregular shaped MCs (lack of granules in part of the cell) following MC depletion, and degranulated cells in response to mCHI. *** p<0.0001, (unpaired, two-tailed Student’s t-test).

We next tested the effect of the prophylactic MC depletion on the development of cephalic pain hypersensitivity following mCHI. A mixed-design ANOVA (**Figure 3B**) revealed time-course decreases in cephalic thresholds (F (3,45) = 22.58; p < 0.0001) that were not affected by the prophylactic MC depletion treatment (F (1,15) = 0.02; p = 0.88). When compared to pre-mCHI baseline values, control (saline-pretreated) animals exhibited decreases in cephalic mechanical pain thresholds at 3 and 7 days post mCHI (q < 0.0001 for 3 and 7 days; post-hoc test). Similarly, when compared to baseline (post 48/80 treatment) MC-depleted animals also exhibited decreased cephalic thresholds at 3, 7, and 14 days (q = 0.002; q < 0.0001; q = 0.03 respectively, post-hoc test). In addition to identifying MC-independent decreases in mechanical thresholds post mCHI, we also observed increases in nociceptive scores (**Figure 3C**) that were time-dependent (F (3,45 = 34.49; p < 0.0001), but not affected by the MC depletion protocol (F(1,15) = 0.3; p = 0.56). In saline-pretreated animals, when compared to pre-mCHI baseline, we identified increases in cephalic scores at 3, and 7 days post mCHI (q < 0.0001; for both, post-hoc test). In MC depleted animals, when compared to pre-mCHI baseline, mCHI gave rise to increased nociceptive scores at 3, 7 and 14 days post mCHI (q < 0.001 for all; post-hoc test). Importantly, analyses of the thresholds and nociceptive scores at baseline, pre-mCHI (following 48/80-induced MC depletion), revealed no differences between the effects of vehicle and 48/80 treatments (q = 0.82 for threshold, q= 0.71 for nociceptive score; post-hoc test), thus confirming that the MC granule depletion protocol *per se* was not pro-nociceptive at the onset of behavioral testing (4 days following the completion of the depletion protocol) and thus unlikely to serve as a confounding variable in the study of mechanical hypersensitivity post mCHI. Taken together, the data suggests that during the acute phase following mCHI induction, functional MCs are not critical for the development of the cephalic pain hypersensitivity.

**Figure 3:**
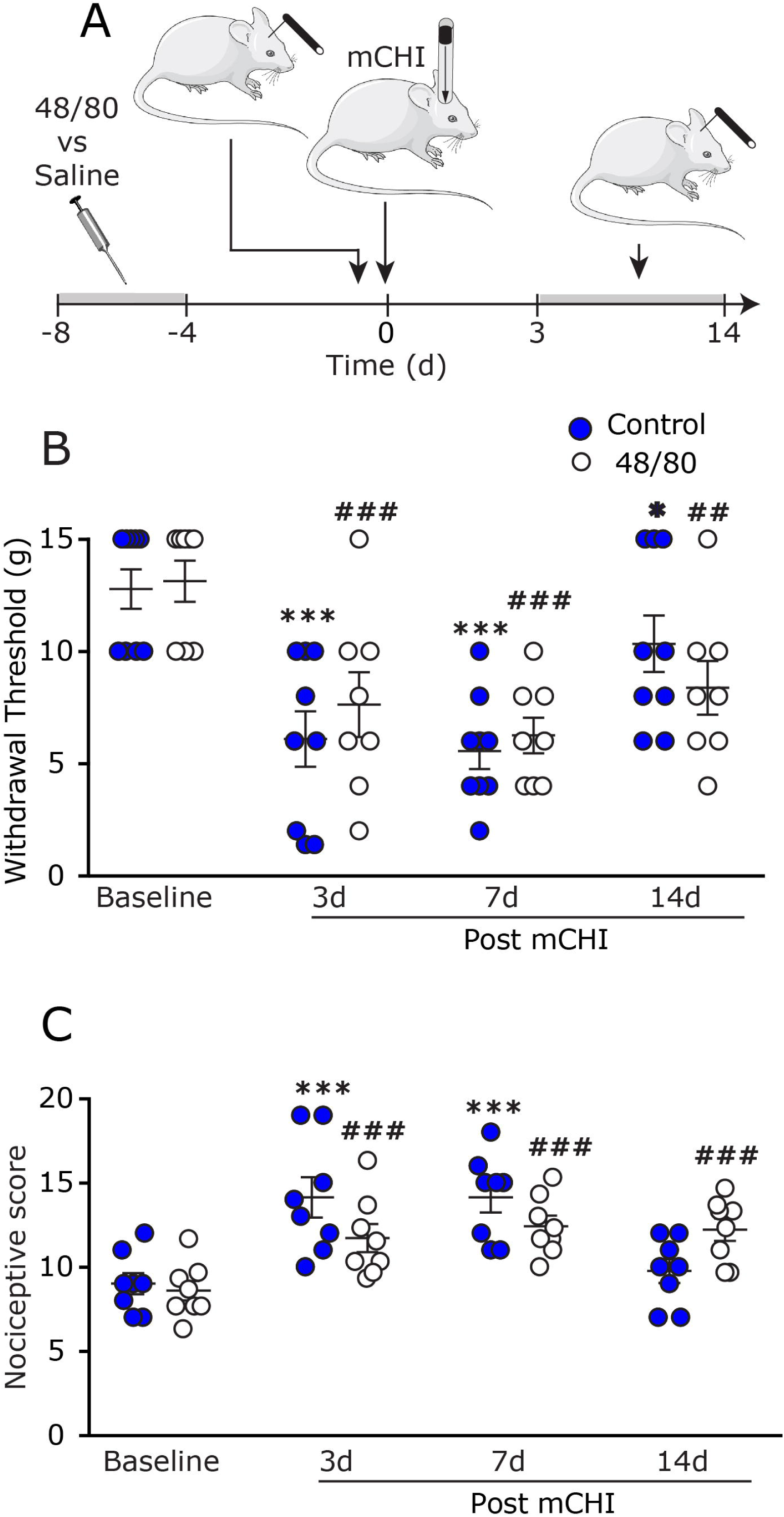
Prophylactic MC depletion does not prevent the development of mechanical pain hypersensitivity following mCHI. (**A)** Scheme of the experimental design. Four days after the end of the MC depletion protocol, or control saline treatment, rats were subjected to baseline behavioral nociceptive testing, followed by mCHI and further behavioral post-mCHI testing, 3-14 days later. Time course changes in cephalic mechanical pain withdrawal thresholds (**B)** and corresponding cumulative nociceptive score (**C)** at baseline, 3, 7, and 14 days post mCHI. *** q < 0.0001 (FDR corrected values following Fisher’s LSD post-hoc test at selected time points vs baseline in control animals). ### q < 0.0001 (FDR corrected values following Fisher’s LSD post-hoc test at selected time points vs baseline in 48/80-treate animals).

### 3.3 Presence of granule-containing MCs during mCHI induction are required for the development of latent mechanical sensitization

We have shown previously that rats subjected to mCHI, but not a sham procedure, develop a prolonged latent mechanical sensitization. This behavior is manifested as decreased cephalic and also extracephalic (hind paw) mechanical pain thresholds in response to systemic administration of a subthreshold dose of the migraine trigger GTN, following the recovery from the acute cephalic hypersensitivity phase [2]. Such latent sensitization has been suggested to play an important role in pain chronification; it likely involves peripheral changes, particularly at the level of the nociceptor’s peripheral nerve endings, and signaling cascades distinct from those underlying acute inflammatory hyperalgesia [37]. We therefore investigated the possibility that, unlike the acute hyperalgesic response to mCHI, the prolonged latent sensitization that develops in the wake of mCHI might involve a MC-related process. Prior to GTN administration, at day 29 following mCHI, baseline cephalic and hind paw mechanical thresholds were not different between animals pretreated with saline or 48/80 prior to mCHI (p = 0.23 and p = 0.88 respectively, unpaired t-test), indicating that MC depletion prior to mCHI induction did not have a long-term effect on mechanical pain responses. In control, saline pre-treated animals, administration of GTN at 30 days post mCHI gave rise to a significant reduction in mechanical pain thresholds in both the cephalic (p < 0.0001, **Figure 4B**, paired t-test) and hind paw (p < 0.0001, paired t-test, **Figure 4D**) regions. However, in animals subjected to the MC granule depletion prior to mCHI, GTN did not lead to a similar pain hypersensitivity. Overall, mechanical withdrawal thresholds tested post GTN treatment were not different than at baseline, in both the cephalic (p = 0.99; **Figure 5C**) and hind paw (p = 0.42; **Figure 4E**) regions. Analyses of the magnitude of GTN-evoked decrease in mechanical pain thresholds also indicated significant differences between the two treatments in both the cephalic region (p = 0.006, unpaired t-test) and hind paw regions (p = 0.04) further indicating that the latent sensitization was blocked by the prophylactic MC depletion.

**Figure 4:**
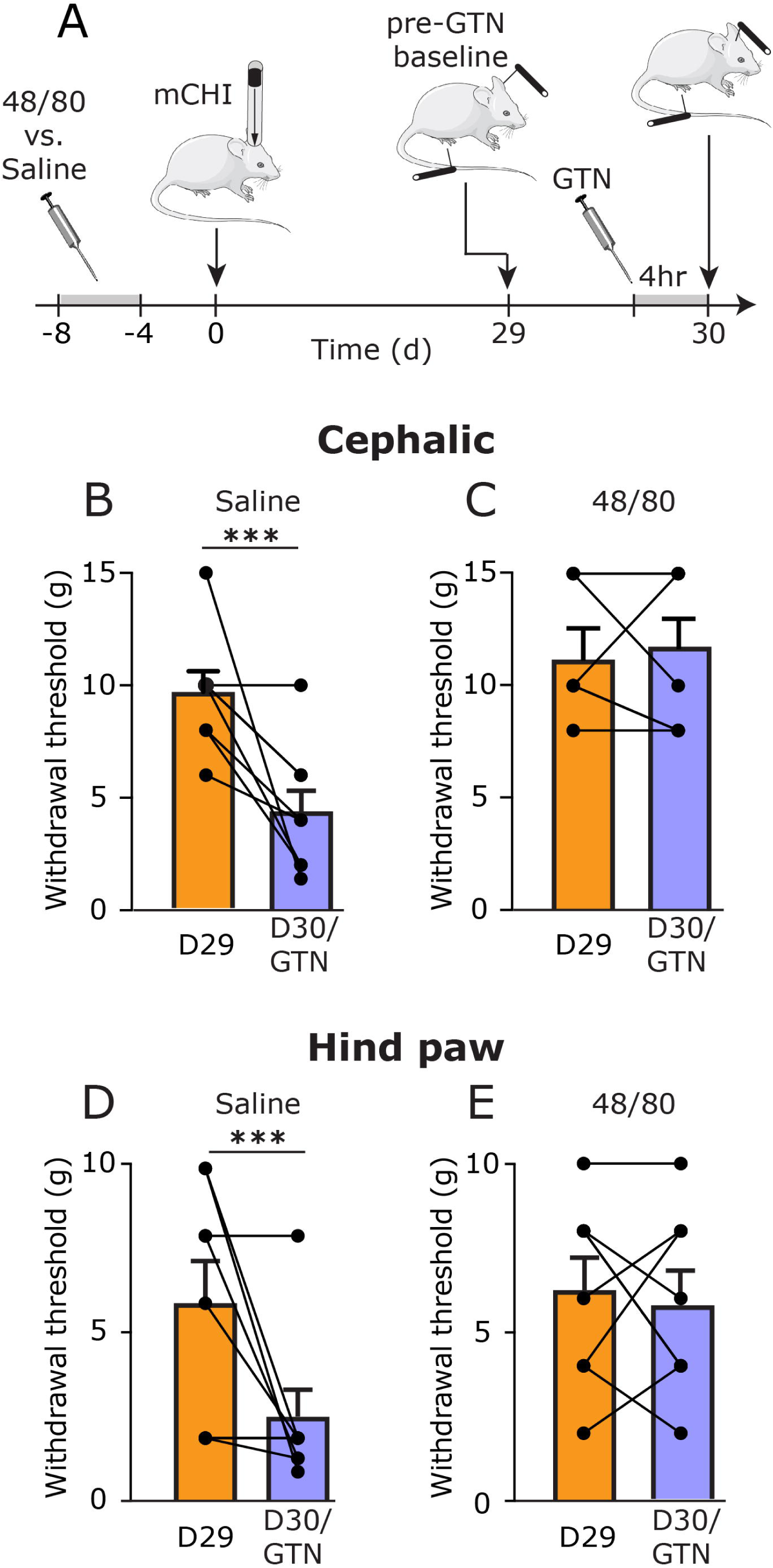
Latent sensitization following mCHI requires an intact MC population during and immediately after the induction of mCHI. (**A)** Scheme of the latent sensitization experimental design. 48/80 and saline-pretreated mCHI rats were subjected to behavioral testing on day 29 following mCHI (pre-GTN baseline), and then a day later (Day 30), 4 hours after GTN administration. Summary of cephalic (**B & C)** and hind paw (**D & E)** mechanical pain thresholds, at baseline (D29; orange bars – mean ± SEM;) and 4 hours following administration of GTN (D30/GTN; purple bars – mean ± SEM;). *** p < 0.0001 pre-GTN baseline vs post GTN (paired student’s t-test).

**Figure 5:**
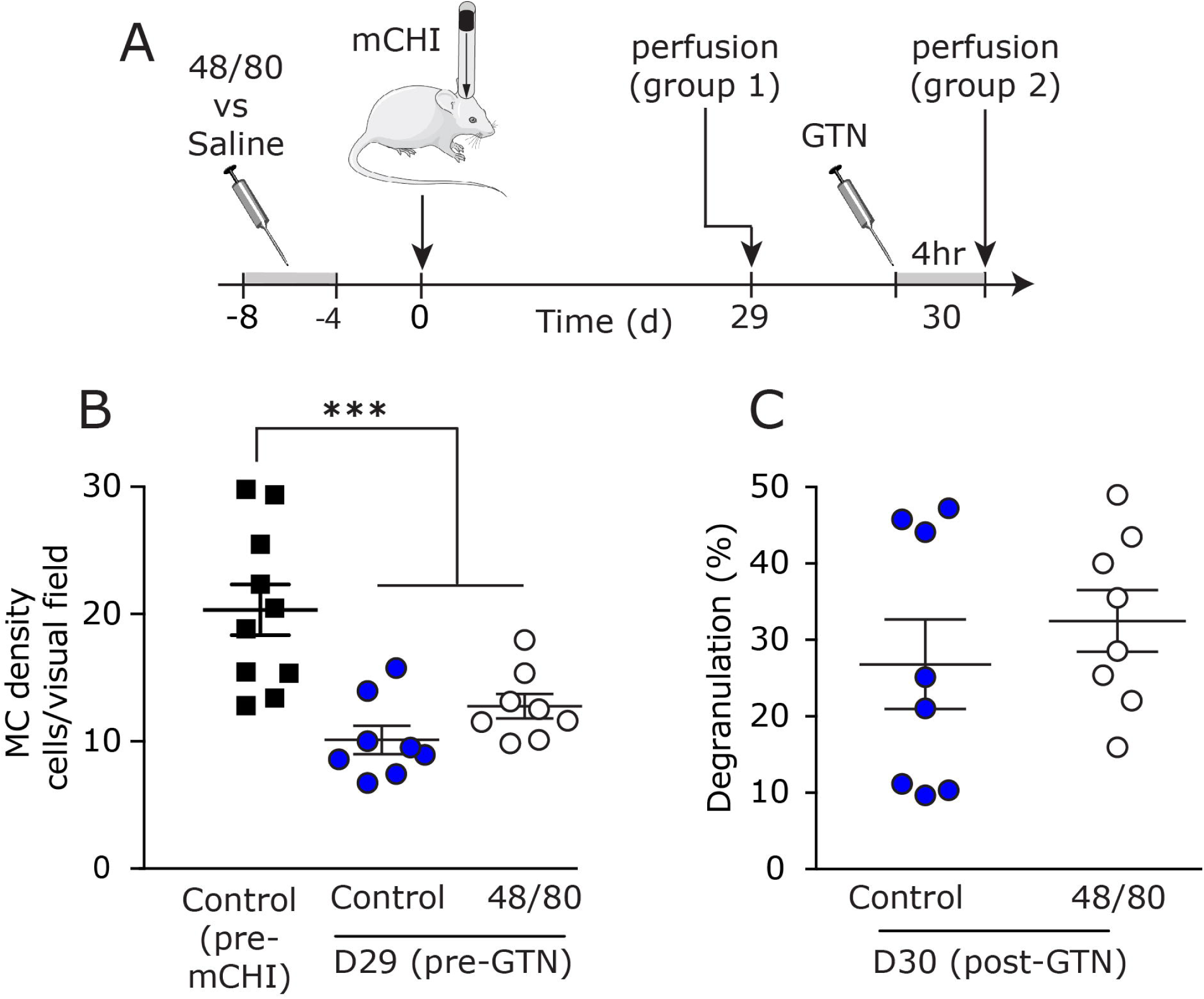
Latent sensitization following mCHI does not involve GTN-evoked acute meningeal MC degranulation. (**A)** Scheme of the experimental design. 48/80 and saline-pretreated mCHI rats were perfused at 29 days post-mCHI or 4 hours after GTN treatment, on day 30 following mCHI. (**B)** Density of meningeal MCs in mCHI animals prior to GTN administration in animals treated prophylactically with 48/80 or saline. (**C)** Meningeal MC degranulation level at 4 hours following GTN treatment in animals pretreated with 48/80 or saline. *** p< 0.001 (unpaired student’s t-test).

### 3.4 Inhibition of the latent sensitization by prophylactic MC granule depletion does not involve altered acute meningeal MC responses to GTN

Systemic administration of GTN has been shown to promote delayed meningeal MC degranulation [36,38]. However, the relative contribution of such proinflammatory response to the activation and sensitization of meningeal afferents and the ensuing behavioral nociceptive effects of GTN in naïve animals is debatable [48]. We therefore tested whether the blockade of the latent sensitization we observed in the 48/80-pretreated mCHI animals might involve altered responses of meningeal MCs, including reduction in the number of cells amenable to degranulation by GTN, as well as their overall level of degranulation following GTN treatment. Prior to GTN administration, the density of non-degranulated meningeal MCs in 48/80-pretereated mCHI animals (34 days following the end of the depletion protocol) was not different than in mCHI animals that received control treatment (saline) prior to mCHI (p = 0.49, unpaired t-test; **Figure 5B**). Both the control (saline-treated) and 48/80-treated groups, however, had lower density of meningeal MCs when compared to pre-mCHI values (p = 0.0003 and p = 0.0053, respectively, unpaired t-test) suggesting that the persistent mCHI-evoked MC degranulation, as seen previously in a mouse mCHI model [18] was not curtailed by the prophylactic MC depletion pre-mCHI. The level of meningeal MC degranulation observed 4 hours following GTN treatment was similar in mCHI animals treated prophylactically with 48/80 and in control, saline-treated animals (p = 0.44, unpaired t-test; **Figure 5C**), suggesting that the inhibitory effect of prophylactic 48/80 treatment protocol on the latent sensitization did not involve a diminished MC degranulating response to GTN.

## 4. Discussion

The main findings of this study suggest that: 1) Ongoing meningeal MC degranulation following mCHI is not ameliorated by treatment with a mAb that blocks peripheral CGRP signaling; 2) The transient cephalic mechanical hypersensitivity that develops following mCHI does not depend upon acute meningeal MC degranulation; 3) Upon the recovery from mCHI-evoked cephalic pain hypersensitivity, the establishment of latent sensitization to GTN requires a MC-related process that likely occurs during, or early after the induction of mCHI, and 4) Latent sensitization to GTN post mCHI does not involve GTN-induced acute meningeal MC degranulation.

The current study, which employed a rat model of mCHI, confirms our previous finding of persistent meningeal MC degranulation in a mouse mCHI model evoked by an analogous weight drop, and suggest a traumatic intracranial injury, with a meningeal inflammatory component. The acute degranulation of meningeal MCs following mCHI likely involves blunt trauma to the calvaria that propagates to the underlying intracranial meningeal tissue [42]. However, the mechanism that maintains this meningeal inflammatory response post-mCHI is unclear. We hypothesized that CGRP, released from activated meningeal afferents, could serve as a key signaling molecule in this process due to its role in promoting meningeal neurogenic inflammation [40], and ability to evoke meningeal MC degranulation [34,39]. However, the current study does not support a role for peripheral CGRP signaling in sustaining the persistent mCHI-evoked meningeal MC degranulation. At present, however, given that the anti-CGRP mAb was administered post-mCHI (to mimic its potential therapeutic use in PTH) we cannot rule out the possibility that CGRP mediates, at least in part, the initial trauma related response of meningeal MCs. Future studies are required to determine whether a prophylactic anti-CGRP mAb treatment (before mCHI) can protect against the initial mCHI-evoked MC degranulation. Whether mCHI leads to persistent activation of meningeal nociceptive afferents, and the ensuing release of other mediators of neurogenic inflammation, such as substance P, or pituitary adenylate cyclase-activating polypeptide (PACAP) that contribute to the prolonged activation of meningeal MCs following mCHI also remains to be studied. The possibility that a non-neurogenic inflammatory process promotes the persistent activation of meningeal MCs following mCHI should also be considered.

We previously hypothesized that the CGRP-dependent cephalic hypersensitivity that develops in the wake of mCHI involves meningeal neurogenic inflammation [2]. The current finding suggesting that persistent activation of meningeal MCs following mCHI is not required for the development of cephalic hypersensitivity indicates, however, that CGRP mediates this PTH-related pain behavior through a mechanism independent of ongoing meningeal MC proinflammatory response. Our data also suggests that the level of meningeal MC degranulation that occurs in the wake of mCHI is not sufficient to promote peripheral meningeal nociception, which likely drives the cephalic pain hypersensitivity after mCHI. While MCs have been implicated in nociceptor sensitization and pain hypersensitivity, some inflammatory conditions have been shown to promote pain hypersensitivity independent of MC activation [22]. At present we cannot discount the possibility that following the depletion protocol the degranulation of a small population of meningeal MCs that underwent partial or complete regranulation prior to the induction mCHI was sufficient to drive the pain behavior following mCHI. However, the earlier finding that the degranulation of up to 50% of the normal population of meningeal MCs (by a headache trigger) is not sufficient to promote the activation of meningeal afferents [38,48] does not support this argument. Whether sterile inflammation propagated by the activation of other meningeal immune cells drives the initial cephalic hypersensitivity in the wake of mCHI remains to be determined. Preliminary data (supplemental Figure 1) suggests that peripheral macrophages also do not play a role in mediating the acute hyperalgesic response following mCHI.

A key, clinically-relevant finding of the rat mCHI model is the development of latent sensitization. This chronic pain related phenomenon is manifested as a delayed and prolonged cephalic and hind paw mechanical allodynia following administration of a subthreshold dose of the headache trigger GTN, long after the resolution of the initial acute cephalic pain hypersensitivity state [2]. A major finding of the current study was that prophylactic depletion of MC granule inhibited the mCHI-evoked latent sensitization. The finding that prior to GTN treatment (at day 29 post mCHI), the density of granulated meningeal MCs was similarly reduced in the control and 48/80-treated animals (presumably as a result of the mCHI) suggests that it was not responsible for the selective lack of the hyperalgesic response to GTN in the 48/80-treated mCHI animals. Our data, showing a similar MC degranulation response to GTN further suggests that the lack of a GTN-evoked hyperalgesic response was not due to reduced acute meningeal MC degranulation, and is in agreement with our previous finding suggesting that the meningeal nociceptive effect of GTN is MC independent [48]. We propose therefore that the MC-related mechanism responsible for mediating the latent sensitization following mCHI involves events that occurred earlier, likely during the acute phase following the mCHI, and that are distinct from those mediating the acute cephalic hyperalgesia. Our previous finding that blocking peripheral CGRP signaling also inhibited the latent sensitization [2], together with our current data suggesting that CGRP does not mediate the mCHI-evoked meningeal MC degranulation, points to the possibility that parallel mechanisms account for the involvement of CGRP and MCs in mediating the latent sensitization. The possibility that this process involves a CGRP-related event that occurs downstream to meningeal MC activation (e.g. NO-mediated signaling [28]) may also be considered.

The latent hyper-responsiveness to GTN, beyond the resolution of the initial acute cephalic hypersensitivity, may be related to a state of hyperalgesic priming - a neuroplasticity response thought to involve persistent and latent hyper-responsiveness of primary afferent nociceptive neurons to inflammatory mediators subsequent to an inflammatory or nerve injury insult [37]. Numerous MC factors have been implicated in the hyperalgesic priming cascade, including the cytokines tumor necrosis factor alpha [35], interleukin-6 [5], nerve growth factor [11], and MC tryptase, via activation of protease-activated receptor 2 (PAR2) [43]. The exact contribution of these mediators and possibly of other MC-related factors, or interaction with other meningeal immune cells, following mCHI will require further studies.

## 5. Conclusions

We evaluated the relative contribution of MCs to the development of acute and persistent pain behaviors reminiscent of PTH following a concussive head trauma in male rats. The key findings point to a differential role of MCs in the genesis and maintenance of PTH: having no contribution to the development of the acute cephalic allodynia, but appearing to be critical for the sustained state of the latent sensitization that develops post-mCHI. Although CGRP has been suggested to play a key role in mediating PTH-like pain behaviors, our data suggests that the mechanism underlying its involvement does not involve downstream signaling mediated by MCs. While further studies are needed to address the role of MCs and their mediators in the genesis of PTH-like symptoms in females, our data suggest that early targeting of MCs and their related peripheral inflammatory and nociceptive signaling following mCHI, at least in males, may be useful to prevent the development of PTH and possibly complementary to treatment with anti-CGRP mAb.

## Supporting information

## Acknowledgment

The authors thanks Dr. Claire (Xinling) Xu (Center for Anesthesia Research Excellence, Beth Israel Medical Deaconess Medical Center, Boston) for advise on the statistical analyses. Authors contributions: Performed experiments: D. Bree; Wrote manuscript: D. Bree and D. Levy; Conceived, designed and lead project; D. Levy.

