## Supplementary material for "Intact mast cell content during mild head injury is required for the development of latent pain sensitization - implications for mechanisms underlying post-traumatic headache"

**Supplemental Figure 1.**

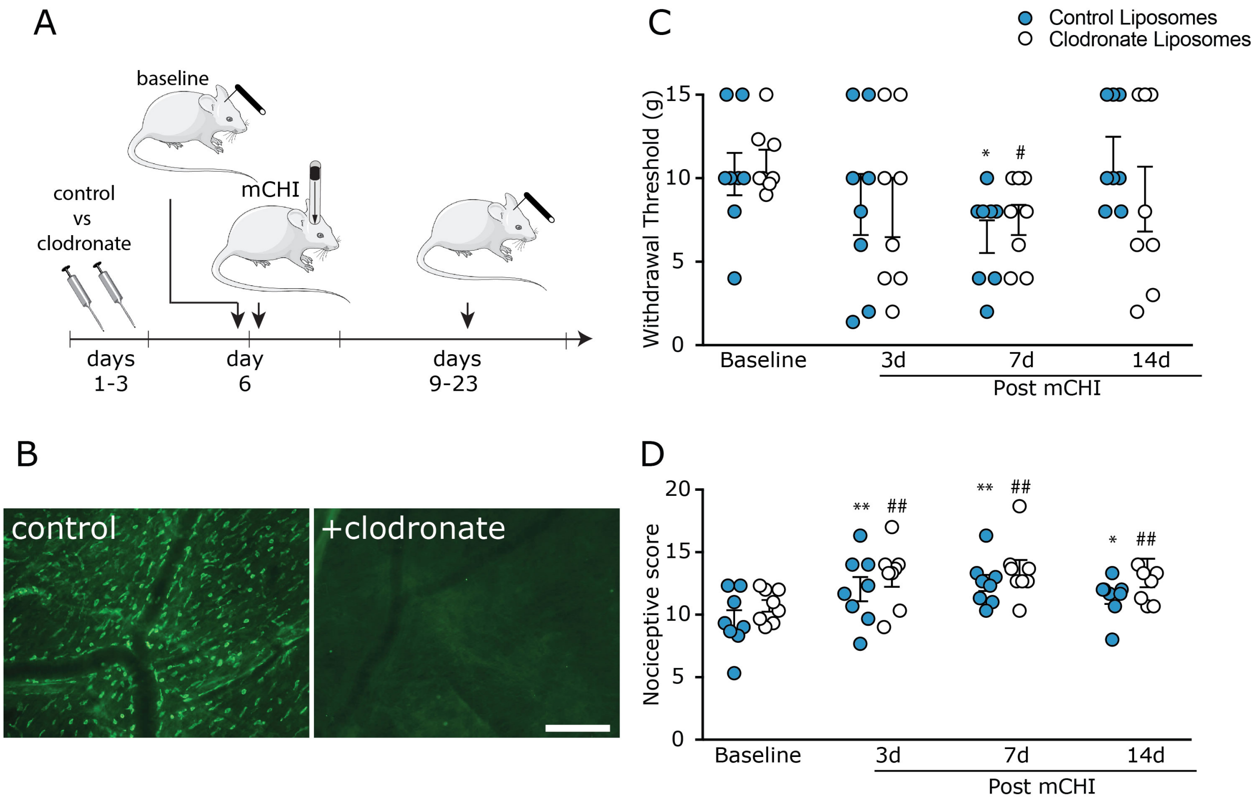


**Supplemental Figure 1. Prophylactic depletion of meningeal macrophages does not prevent the development of mechanical pain hypersensitivity following mCHI**. (A) Rats were subjected to a macrophages depletion using i.p. injections of clodronate liposomes, or control liposomes (https://clodronateliposomes.com) at 1ml/kg. The concentration of clodronate is 5mg/kg. Two injections were made, 48 hr apart, and rats were subjected to baseline behavioral testing 72 hrs later followed by isoflurane anesthesia and induction of mCHI. Post-mCHI behavioral testing were conducted 3-14 days later. (B) Representative images of meningeal whole-mounts subjected to immunohistochemistry using a mouse anti-rat mAb against CD163 (clone ED2, 1:500, MCA342, BioRad) showing dural macrophages in animals treated with control liposomes and their depletion following treatment with clodronate. Scale bar = 500µm. Cephalic mechanical pain withdrawal thresholds (C) and corresponding cumulative pain response scores (D) at baseline, 3, 7, and 14 days post mCHI. Macrophage depletion did not have an effect on the mCHI-evoked decrease in cephalic mechanical pain threshold $(F_{1,14}) = 0.03$; $p = 0.86$), or the increase in pain score in response to mechanical stimulation $(F_{1,14}) = 1.9$; $p = 0.19$). * $q < 0.05$, ** $q < 0.01$ control liposomes vs pre-mCHI baseline; # $q < 0.05$, ## $q < 0.01$ liposomes vs pre-mCHI baseline.
